# A snapshot of statistical methods used in experimental immunoblotting: A scoping review

**DOI:** 10.1101/2022.04.06.487284

**Authors:** Romain-Daniel Gosselin

## Abstract

Among the many avenues considered to make life science more reproducible, the improvement of the quality and openness of statistical methods has taken centre stage. However, although disparities across research fields and techniques are not unknown, they remain largely unexplored. The present scoping review aims to provide a first overview of statistical methods used in biochemical research involving immunoblotting (also referred to as western blotting), a technique frequently used to quantify proteins of interest. A reverse chronological systematic sampling was implemented to explore the PubMed online browser and analyse 2932 experimental conditions (i.e., experimental groups) in quantitative immunoblots from 64 articles published at the end of 2021. The statistical test (actual study size n = 67) and software (actual study size n = 61) used for each article and the sample size for each experimental condition were documented. The results indicate an omnipresence of parametric tests, mostly one-way analysis of variance (ANOVA, 15/67) and Student’s t-test (13/67), but for many articles the statistical procedure was not clearly divulged (23/67). GraphPad Prism was the most commonly used statistical package (36/61), but many (14/61) articles did not reveal the package used. Finally, the sample size was clearly disclosed in only 1054/2932 conditions in which its median value was 3 (IQR = [3 - 6]). This study suggests that the transparency of reporting might be suboptimal in immunoblotting research and prompts more comprehensive systematic reviews in future.

## Introduction

Transparency, correctness, and quality in published protocols are key prerequisites to ensure replication in science and enable evidence-based research practice. In particular, the use of inappropriate statistical designs and methods as well as their insufficient reporting are considered important contributors to the so-called reproducibility crisis in life science [1]. Guidelines describing statistical reporting are regularly released, thereby establishing an important framework of standardization for published material. These include Animal Research: Reporting of In Vivo Experiments (ARRIVE, recently updated as ARRIVE 2.0) [2, 3], the Checklist for Reporting In-vitro Studies (CRIS) [4], guidelines from the American Physiological Society [5] and the checklist by Emmerich and Harris for in vitro research [6]. Although the aforementioned guidelines are generally addressed to a well-defined scientific community, the current statistical practices in reporting across different research domains remain largely unknown. In this context, meta-research that describes the characteristics of statistical methods in specific research fields would ultimately enable the implementation of targeted educational initiatives in defined research communities.

Immunoblotting (also referred to as western blotting) is a very common technique in biochemistry that uses protein separation by molecular weight using polyacrylamide gel electrophoresis (SDS-PAGE) and a subsequent antibody-based specific detection and relative quantification of proteins of interest. Others have explained that method disclosure in research based on antibodies, including immunoblotting, was not sufficient to enable reproducibility, which prompted the release of specific reporting guidelines in the field [7, 8]. However, these recommendations focused primarily on the disclosure of reagents, laboratory protocols and methods of image acquisition/processing with only minor sections on statistical analysis of quantitative blots. The present scoping review aims to assess statistical methods used in immunoblotting to potentially rationalize further actions such as comprehensive systematic reviews or the assembling of specific guidelines. The results obtained in a large sample of articles suggest that statistical design and reporting are of an insufficient standard in immunoblotting research.

## Material and Methods

This scoping review was designed and prepared according to the 2018 Preferred Reporting Items for Systematic reviews and Meta-Analyses extension for Scoping Reviews (PRISMA-ScR)[9] and the 2020 “Updated methodological guidance for the conduct of scoping reviews” [10, 11]. The dual indication was to “examine how research is conducted on a certain topic or field” and serve “as a precursor to a systematic review”. In addition, this manuscript was also prepared in accordance with the ACcess to Transparent Statistics (ACTS) guideline [12]. Therefore, the methods section starts with the statistical paragraph that includes the eight components of ACTS and the discussion incorporates a specific section dedicated to the limitations of the study. For the sake of clarity, throughout the text, the term “sample size” refers to the number of experimental conditions in immunoblots in the articles included in the scoping review, and the term “study size” refers to the number of analytical units in the present scoping review (either the number of articles or total number of experimental conditions included). The study protocol was not preregistered.

## 1 Statistical methods

Data were collected, structured, and managed using Microsoft Excel for Mac (version 16). Calculations of descriptive statistics (quartiles, interquartile range) and the drawing of bar charts (Figure 2) and the histogram (Figure 3) were performed using the base function in R software (R-studio, version 4.1.1). The statistical items quantified in the study were: the software used, the procedures (tests) implemented, and the number of observations collected (i.e., sample size). The analytical units for the documentation of software and tests were the articles (i.e., the items were listed for each article). For the assessment of sample size, the analytical units were the separate observations (bands), presumed independent, used by the authors in each condition. The study size was a priori determined using a precision-based approach applying the following formula:

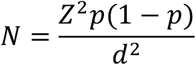

N is the minimum number of articles required to establish a 95% confidence interval (CI, Z=0.95) for software and tests if the least frequent software or test has a frequency (p) of 0.1 (i.e., present in 10% of articles), with a precision (d) of 0.07 (i.e., 7%). The calculation gave a minimum size of 70 articles for the study.

## 2 Article sampling

Articles with quantitative western blots were sampled from the PubMed browser of the NCBI website (https://pubmed.ncbi.nlm.nih.gov/) using reverse chronological convenience sampling (Figure 1, Table 1). In the advanced NCBI research tool, the following terms were entered for query: “immunoblot” OR “western blot” OR “western-blot” OR “sds page” OR “sds-page” and the NCBI publication filter was used with “end date” set at 31.12.2021. Retrieved articles were individually checked for compliance with inclusion criteria and absence of exclusion criteria by clicking on the link to the journal page or DOI until 70 articles were collected. In particular, the article text and figures were quickly examined to confirm both that the authors performed an immunoblot (and not a mere polyacrylamide gel electrophoresis) and that at least one reported blot was quantitative (presence of statistics and/or of text providing a quantitative interpretation of the blots). Twenty-six publications were excluded at this stage. During the subsequent detailed analysis of statistical features (see infra), six additional articles were excluded because they did not perform quantitative immunoblots, contrary to what was presumed upon first reading. The final list of included (n = 64) and excluded articles is provided on the Figshare public repository (https://doi.org/10.6084/m9.figshare.19448264.v1).

**Figure 1:**
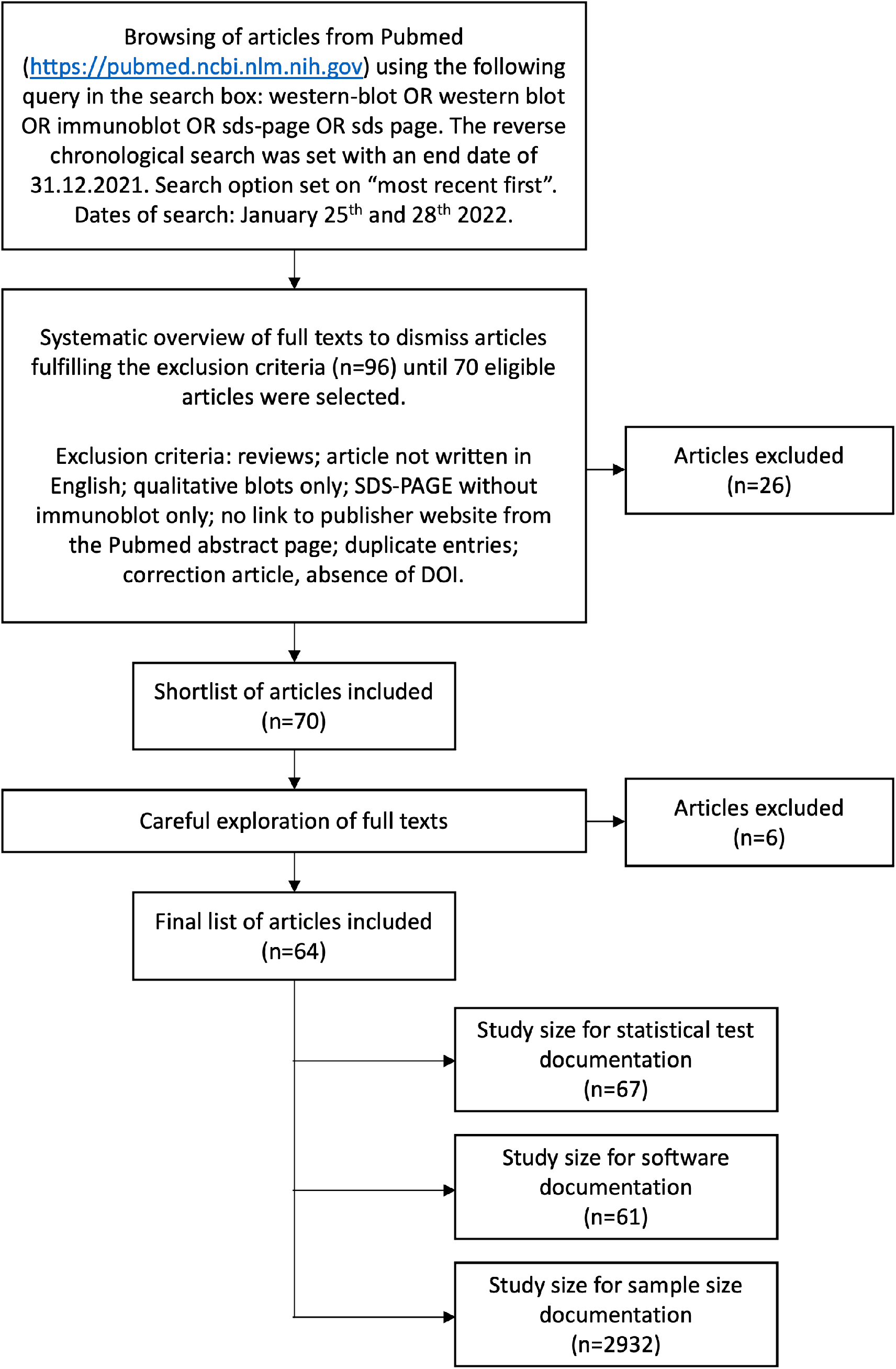
Flow chart of the sampling protocol.

**Table 1:**
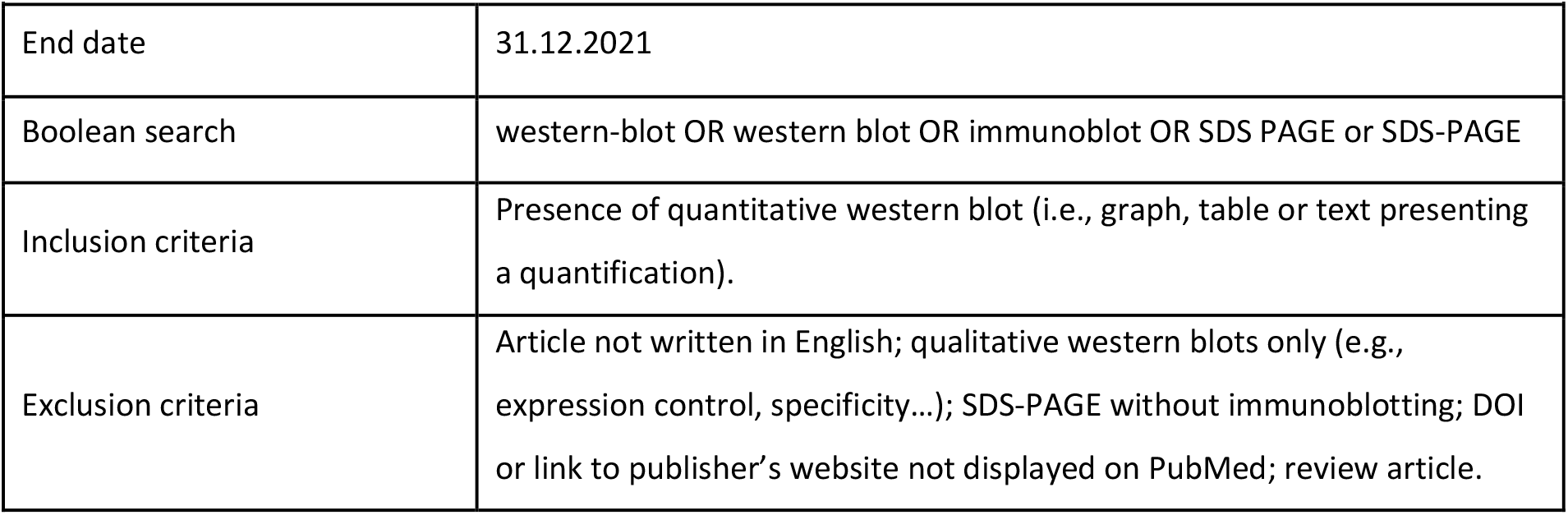
Selection criteria applied to the PubMed search engine

Out of the 64 sampled articles, two articles used multiple procedures to analyse their immunoblots. One publication used both Student’s t-test and an unknown post-test after an ANOVA, and no statistical analysis for a blot interpreted quantitatively. Another article used both an unclear test for one immunoblot figure and no statistical analysis for a western blot interpreted quantitatively. Therefore, the final study size used in the present description of statistical tests used is n = 67. On the other hand, three sampled articles did not provide any statistical analysis or statistical software although they reported quantitative analyses, therefore the final study size for the documentation of software is n = 61 (Figure 1).

## 3 Quantification of statistical features

For each article sampled, for the experiments that reported quantitative immunoblots, the corresponding text (methods, results, discussion, figure legends) was explored, and three statistical items were documented: the statistical software used, the statistical procedure (e.g. test) utilized and the sample size (disclosed number of observations) in each experimental condition. Immunoblots in figures provided as supplemental materials were not included, but supplemental methods referring to figures in the main text were considered full-fledged methodological information. The category “not disclosed/unclear” corresponds to instances where the item was not given, although a statistical analysis (e.g., p-value, discussion on statistical significance) was provided, or was not given in an unambiguous manner. Absence of clear information about a post-test used after ANOVA was included in this latter category.

## 4 Data and code availability

All data generated and analysed during this study, as well as the R code used, are publicly available on Figshare at https://doi.org/10.6084/m9.figshare.19448264.v1.

## Results

The descriptive analysis of statistical procedures is presented in Figure 2a. The results revealed widespread use of parametric tests, namely one-way analysis of variance (ANOVA) with post-test (15/67), Student’s t-test (regardless of their laterality or paired vs. unpaired nature, 13/67) and two-way ANOVA with post-test (5/67). Non-parametric tests were almost absent, with only Wilcoxon signed-rank test (1/67), Kruskal–Wallis test (followed by a post-test 1/67), and Duncan’s multiple range test (1/67). Strikingly, in a large proportion of quantitative analyses, the statistical procedure was unclear or not disclosed (23/67, which included instances where the post-test was not disclosed after the omnibus procedure), or no statistical analyses were performed (8/67). The evaluation of statistical software shows that only three packages were cited (Figure 2b). The most frequently used was GraphPad Prism (36/61), followed by IBM SPSS (10/61) and the open-source software R (1/61). Fourteen articles (14/61) did not cite any software.

**Figure 2:**
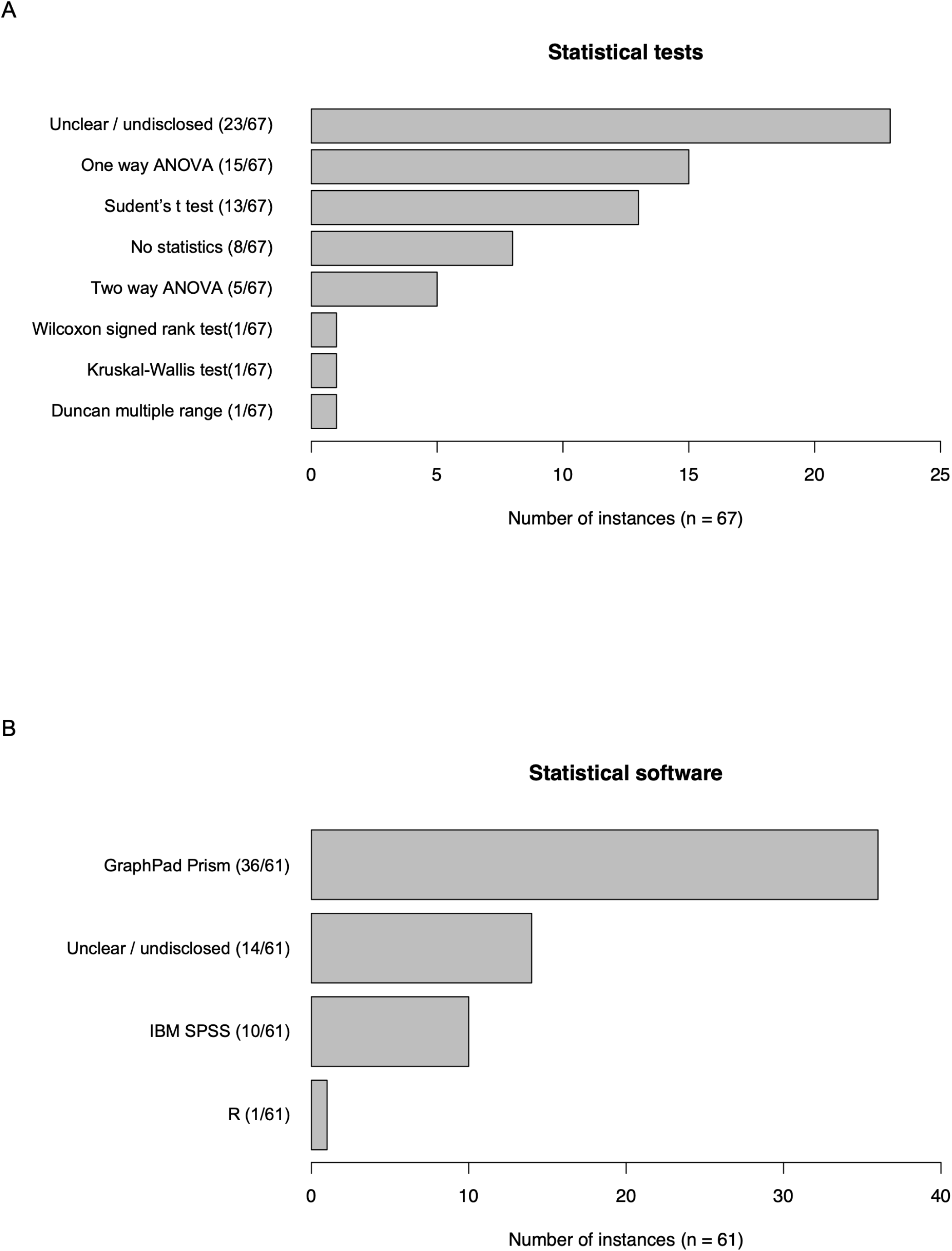
Quantification of statistical methods (a, study size: n = 67) and software (b, study size: n = 61) in articles (n = 64) with quantitative immunoblotting.

The sample size was clearly indicated in only 1054/2932 experimental conditions and was conversely often unclear (e.g., when inequalities were given instead of exact numbers) or not disclosed (1592/2932). The frequency distribution of sample sizes (Figure 3) shows the vast over-representation of very small samples, with 744/1054 experimental conditions that incorporated fewer than four observations per sample. In about 10% of experimental conditions (286/2932), the notion of sample size was even irrelevant due to the use of single blots to quantitatively interpret the experiment. The analysis of disclosed sample sizes shows that the median number of observations (i.e., individual blots) used in quantitative immunoblots is 3 (range [2–40], interquartile range, IQR [3 - 6]).

**Figure 3:**
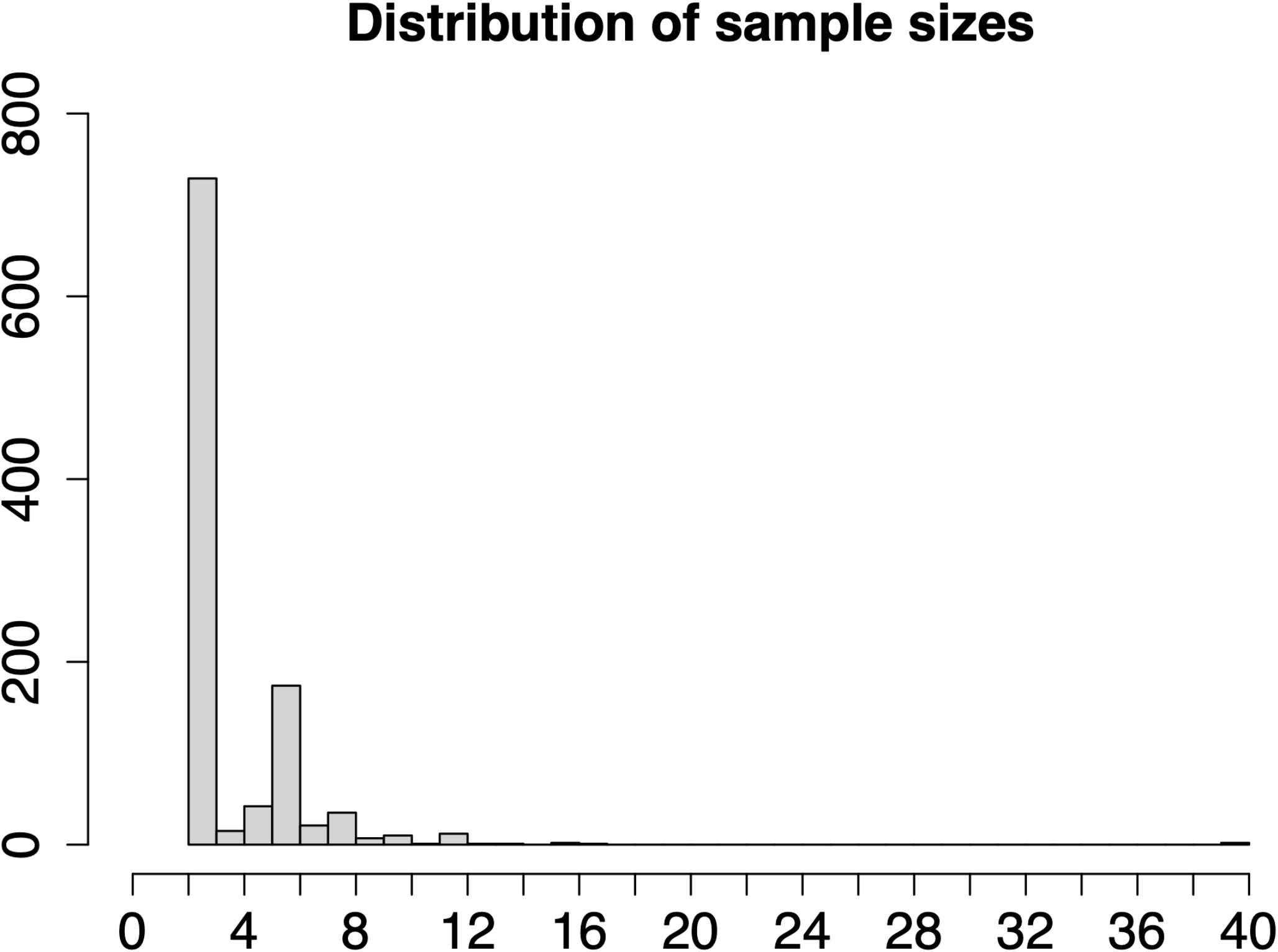
Histogram showing the frequency distributions of sample size in immunoblotting experimental conditions (actual study size: n = 1054). Only experimental conditions with clearly disclosed sample sizes were included (1054/2932).

## Discussion

In the sample analysed in the present scoping review, the quantitative immunoblotting was typically designed with small numbers of observations and analysed with parametric tests using proprietary software. The use of such designs may jeopardize their validity and sensitivity (i.e., statistical power). First, the verification of parametric assumptions, which are a mandatory prerequisite for the implementation of parametric tests, is challenging when samples are small. Indeed, in that context, graphical methods such as quantile-quantile (Q-Q) plots are hardly reliable and distribution tests for normality (e.g. Shapiro-Wilk, Kolmogorov-Smirnov) tend to be largely underpowered to detect any deviation from normality. Furthermore, statistically significant differences obtained with small samples tend to overestimate effect sizes and to have a higher percentage of false positives among statistically significant results [13]. In addition, small sample sizes such as observed herein tend also to have low statistical power and thus inflate the fraction of false negative results. This shrinkage of statistical power may even engender an impossibility to detect statistically significant differences at the usual confidence levels if non-parametric tests are used [14]. Therefore, for all these reasons, the sampled literature that reports quantitative immunoblotting might be riddled with a large share of statistically unreliable results.

In the present sample, a large proportion of statistical methods is not disclosed, data that is in line with the lack of transparency recurrently reported in other studies [15-17], which is a strong impediment to reproducibility. Many guidelines exist but their publication and dissemination has shown only limited impact. Therefore, although this route of action appears necessary, especially to provide a framework of official recommendations, it is probably not sufficient to initiate a cultural change towards a more reproducible culture. Strengthening continuing education programmes on good research practices for graduate students, post-doctoral researchers and established researchers would be an important avenue to explore. The organisation of such education for professionals at career stages where improving methodological and statistical literacy relates to the actual daily laboratory activity might be more efficient than targeting young students at the undergraduate level. Furthermore, in methodological courses, a marked emphasis should be placed on experimental design and data sampling rather than mere data analysis and statistical inference [18].

The question of whether research involving immunoblotting uses different standards of design, analytic methods and reporting than other fields of experimental science would be unequivocally answered by future comprehensive systematic reviews. Should it be the case, measures assembled to particularly target this research community would be warranted, including guidelines in field-specific journals, thematic sessions on design and transparency in scientific events or more active education on these topics in biochemistry curricula.

Importantly, the alarmingly high prevalence of non-transparent methods points also to a similar insufficient statistical literacy among peer reviewers in the field. Although guidelines articles intended for peer reviewers are available [7, 19], peer reviewing and editorial filters do not apparently constitute efficient bottlenecks for statistical quality. One solution might be the systematic invitation of statistical reviewers to help ensure standards in statistical reporting while, at the same time, lifting some of the reviewing burden from non-statistical reviewers [20]. Nevertheless, it is conceivable to restrict this option to manuscripts with relatively complex designs, leaving non-statistical reviewers to perform an assessment of statistical transparency and elementary statistics in most studies. In this respect, standardization of minimal statistical knowledge among reviewers is important and would benefit from the widespread creation of continuing education programmes in manuscript peer reviewing [21]. Authoring and reviewing activities are fulfilled by overlapping groups of scholars, and the concepts to be assimilated are the same. Therefore, it is also suggested here that guidelines on reporting practices should systematically be clearly addressed to both researchers and peer reviewers and make explicit that better knowledge of such good practices is equally important when engaging in either activity. In addition, specific educational programmes on biostatistics and reporting organized in institutes that perform immunoblotting research should similarly unambiguously clarify that the exposed concepts apply similarly to the design, analysis, and presentation of a research project and its assessment as a reviewer.

In conclusion, the results indicate that small samples, parametric tests and proprietary software might be omnipresent in immunoblotting research and that non-transparency in statistical protocols is widespread. Targeting researchers in this specific field with continuing education on good research practice and fundamental biostatistics should be a priority objective. Additional avenues to explore are better communication or enforcement of existing reporting guidelines. Journals and editors should take a leading role in these efforts.

## Limitations

The present scoping review has several limitations. First, other statistical features could have been considered to achieve a more comprehensive description of statistical reporting. These include the choice of error, the nature of post-tests, or the significance thresholds used. In addition, this study was designed as descriptive and does not intend to make inferences about all of the literature that uses western blots. For example, the date of publication selected and the convenience sampling method may have created an over-representation of some journals and editors that prevents generalization. Nevertheless, the consistency of the results within this sample is already notable, suggesting a widespread entrenched culture of incomplete reporting in immunoblot research and prompts for future confirmatory systematic review. Furthermore, the study protocol was not preregistered, which constitutes another limitation. Finally, the present study has been designed and analysed by one single reviewer, a methodology deemed acceptable for accelerated evidence synthesis such as the current scoping reviews [22, 23] but which might increase error and bias for comprehensive systematic reviews [23, 24].

## Acknowledgement

The author would like to thank Prof Jacques Fellay for his support. The research was supported by institutional intramural funding from Lausanne University Hospital (CHUV).

## Author contributions

RDG designed the study, analysed the data, and wrote the manuscript.

---

Preferred Reporting Items for Systematic reviews and Meta-Analyses extension for Scoping Reviews (PRISMA-ScR) Checklist

| SECTION | ITEM | PRISMA-ScR CHECKLIST ITEM | REPORTED ON PAGE # |
| --- | --- | --- | --- |
| **TITLE** |  |  |  |
| Title | 1 | Identify the report as a scoping review. | 1 |
| **ABSTRACT** |  |  |  |
| Structured summary | 2 | Provide a structured summary that includes (as applicable): background, objectives, eligibility criteria, sources of evidence, charting methods, results, and conclusions that relate to the review questions and objectives. | 2 |
| **INTRODUCTION** |  |  |  |
| Rationale | 3 | Describe the rationale for the review in the context of what is already known. Explain why the review questions/objectives lend themselves to a scoping review approach. | 3 |
| Objectives | 4 | Provide an explicit statement of the questions and objectives being addressed with reference to their key elements (e.g., population or participants, concepts, and context) or other relevant key elements used to conceptualize the review questions and/or objectives. | 2, 3 |
| **METHODS** |  |  |  |
| Protocol and registration | 5 | Indicate whether a review protocol exists; state if and where it can be accessed (e.g., a Web address); and if available, provide registration information, including the registration number. | 4 |
| Eligibility criteria | 6 | Specify characteristics of the sources of evidence used as eligibility criteria (e.g., years considered, language, and publication status), and provide a rationale. | 4, 5, figure 1, table 1 |
| Information sources* | 7 | Describe all information sources in the search (e.g., databases with dates of coverage and contact with authors to identify additional sources), as well as the date the most recent search was executed. | 4 |
| Search | 8 | Present the full electronic search strategy for at least 1 database, including any limits used, such that it could be repeated. | 4, 5, figure 1, table 1 |
| Selection of sources of evidence† | 9 | State the process for selecting sources of evidence (i.e., screening and eligibility) included in the scoping review. | 4, 5, figure 1, table 1 |
| Data charting process‡ | 10 | Describe the methods of charting data from the included sources of evidence (e.g., calibrated forms or forms that have been tested by the team before their use, and whether data charting was done independently or in duplicate) and any processes for obtaining and confirming data from investigators. | 4, 5, figure 1, table 1 |
| Data items | 11 | List and define all variables for which data were sought and any assumptions and simplifications made. | 5 |
| Critical appraisal of individual sources of evidence§ | 12 | If done, provide a rationale for conducting a critical appraisal of included sources of evidence; describe the methods used and how this information was used in any data synthesis (if appropriate). | 4, 5, 8, 9 |
| Synthesis of results | 13 | Describe the methods of handling and summarizing the data that were charted. | 4, 5 |
| **RESULTS** |  |  |  |
| Selection of sources of evidence | 14 | Give numbers of sources of evidence screened, assessed for eligibility, and included in the review, with reasons for exclusions at each stage, ideally using a flow diagram. | 4, 5, figure 1, data provided (excel file) |
| Characteristics of sources of evidence | 15 | For each source of evidence, present characteristics for which data were charted and provide the citations. | data provided (excel file) |
| Critical appraisal within sources of evidence | 16 | If done, present data on critical appraisal of included sources of evidence (see item 12). |  |
| Results of individual sources of evidence | 17 | For each included source of evidence, present the relevant data that were charted that relate to the review questions and objectives. | data provided (excel file) |
| Synthesis of results | 18 | Summarize and/or present the charting results as they relate to the review questions and objectives. | 6, fig 2, fig3 |
| **DISCUSSION** |  |  |  |
| Summary of evidence | 19 | Summarize the main results (including an overview of concepts, themes, and types of evidence available), link to the review questions and objectives, and consider the relevance to key groups. | 6, 7, 8 |
| Limitations | 20 | Discuss the limitations of the scoping review process. | 8 |
| Conclusions | 21 | Provide a general interpretation of the results with respect to the review questions and objectives, as well as potential implications and/or next steps. | 8 |
| **FUNDING** |  |  |  |
| Funding | 22 | Describe sources of funding for the included sources of evidence, as well as sources of funding for the scoping review. Describe the role of the funders of the scoping review. | 1, 9 |

JB1 = Joanna Briggs Institute; PRISMA-ScR = Preferred Reporting Items for Systematic reviews and Meta-Analyses extension for Scoping Reviews.
\* Where *sources of evidence* (see second footnote) are compiled from, such as bibliographic databases, social media platforms, and Web sites.
† A more inclusive/heterogeneous term used to account for the different types of evidence or data sources (e.g., quantitative and/or qualitative research, expert opinion, and policy documents) that may be eligible in a scoping review as opposed to only studies. This is not to be confused with *information sources* (see first footnote).
‡ The frameworks by Arksey and O'Malley (6) and Levac and colleagues (7) and the JBI guidance (4, 5) refer to the process of data extraction in a scoping review as data charting.
§ The process of systematically examining research evidence to assess its validity, results, and relevance before using it to inform a decision. This term is used for items 12 and 19 instead of "risk of bias" (which is more applicable to systematic reviews of interventions) to include and acknowledge the various sources of evidence that may be used in a scoping review (e.g., quantitative and/or qualitative research, expert opinion, and policy document).
From: Tricco AC, Lillie E, Zarin W, O'Brien KK, Colquhoun H, Levac D, et al. PRISMA Extension for Scoping Reviews (PRISMA-ScR): Checklist and Explanation. *Ann Intern Med*. 2018;169:467–473. doi: 10.7326/M18-0850.

